# Microevolutionary Hypothesis of the Obesity Epidemic

**DOI:** 10.1101/2023.08.29.555238

**Authors:** Joseph Fraiman, Scott Baver, Maciej Henneberg

## Abstract

The obesity epidemic represents potentially the largest phenotypic change in *Homo sapiens* since the origin of the species. Despite obesity’s high heritability, a change in the gene pool has not generally been presumed as a potential cause of the obesity epidemic. Here we advance the hypothesis that a rapid change in the obesogenic gene pool has occurred second to the introduction of modern obstetrics dramatically altering evolutionary pressures on obesity - the microevolutionary hypothesis of the obesity epidemic. Obesity is known to increase childbirth related mortality several fold. Prior to modern obstetrics, childbirth related mortality occurred in over 10% of women. After modern obstetrics, this mortality reduced to a fraction of a percent, thereby lifting a strong negative selection pressure. Regression analysis of data for ∼ 190 countries was carried out to examine associations between 1990 maternal death rates (MDR) and current obesity rates. Multivariate regression showed MDR correlated more strongly with national obesity rates than GDP, calorie intake and physical inactivity. Analyses controlling for confounders via partial correlation show that MDR explains approximately 11% of the variability of obesity rate between nations. For nations with MDR above the median (>0.45%), MDR explains over 20% of obesity variance, while calorie intake, and physical inactivity show no association with obesity in these nations. The microevolutionary hypothesis offers a parsimonious explanation of the global nature of the obesity epidemic.

**Significance Statement:** Humans underwent a rapid increase in obesity in the 20^th^ century, and existing explanations for this trend are unsatisfactory. Here we present evidence that increases in obesity may be in large part attributable to microevolutionary changes brought about by dramatic reduction of childbirth mortality with the introduction of modern obstetrics. Given the higher relative risk of childbirth in women with obesity, obstetrics removed a strong negative selection pressure against obesity. This alteration would result in a rapid population-wide rise in obesity-promoting alleles. A cross-country analysis of earlier maternal death rates and obesity rate today found strong evidence supporting this hypothesis. These findings suggest recent medical intervention influenced the course of human evolution more profoundly than previously realized.

## Introduction

Over the last half-century, the world has been experiencing a pandemic less immediately dangerous than the current COVID one, but more insidious - obesity. The increasing rate of obesity appeared to begin in developed countries, but it is now a global health crisis. The diagnostic tool commonly used to detect obesity is Body Mass Index which, despite being fraught with some inadequacies, gives a general picture of increasing adiposity. Obesity, as a pathological state, consists of excessive accumulation of white adipose tissue, not just increase in body mass. It is commonly accepted in the scientific literature, medical community, and population at large that positively altered energy balance - more energy consumed than energy expended - leads to accumulation of adipose tissue in the body. This accumulation, however, varies among individuals and may be dependent on body build and physiology. This notwithstanding, a vast quantity of research has been made on the regulation of energy balance and multiple dietary and exercise interventions have been introduced over the last decades to reduce obesity. Alas, to no, or little, avail, since the prevalence of obesity has increased in all nations.

Obesity is a highly heritable trait with twin studies finding heritability between 74% and 90% when only including studies in which zygosity of twins is confirmed (1). Yet it is commonly stated that a rapid change in the gene pool can not have caused the obesity epidemic in such a short period of time (2). Given this reasoned out conclusion, a focus on the genetic basis to the obesity epidemic has been limited. Here we present the hypothesis that there has been a ‘rapid’ change in the gene pool as an explanation for an underlying rationale for the obesity epidemic. We term this the “microevolutionary hypothesis of the obesity epidemic” . Prior to the introduction of modern obstetrics, childbirth deaths of mother and/or child were commonplace and well known to have placed a strong negative selective pressure shaping the course of human evolution. With the introduction of modern obstetrics in the 1930s, childbirth deaths reduced dramatically (3, 4), to a level negligible as an evolutionary pressure. Obesity is a well known risk factor for maternal and perinatal mortality with the relative risk of death being several fold higher for obese women (5). This increased relative risk of obesity placed on childbirth without obstetric care would also be a high absolute risk exerting a strong negative selection pressure against obesity. Despite the fact that childbirth exerts a strong selection pressure against obesity, all human populations retain obesity promoting genes suggesting the existence of a stabilizing component of selection pressure against increased body mass combined with stabilizing pressures against genes causing reductions in body mass such as, for instance anorexia, to maintain “normal” body mass. Modern obstetrics removed the strong selection pressure by childbirth. This would, in a few generations, cause a rapid increase in obesogenic genes in the population. An imbalance in stabilizing selection pressures on obesity-related genes has occurred producing a directional change towards obesity.

### Is a Rapid Change in the Gene Pool Possible?

The widespread belief that the obesity epidemic could not be caused by a rapid change in the gene pool developed prior to the understanding that rapid changes in the gene pool can and do occur. It is now known that these rapid changes occur in a population following a sudden environmental change that dramatically alters evolutionary pressures on a trait with large-standing genetic variation (6). Obesity is known to have large standing genetic variation, (with over 1,000 associated obesity promoting loci (7) each with near negligible phenotypical penetrance of each allele. This has led to the problem in genome-wide association studies (GWAS) of BMI only identifying about 3% of the alleles of this large standing variation (8). This phenomenon is referred to as hidden heritability, and it occurs when GWAS studies can identify only a small percentage of the genes controlling a highly heritable phenotype (9). To understand how this rapid change in gene pool could occur with large standing variation consider if there are 1,000 obesity associated loci and if, hypothetically, each allele increases an individual’s BMI by only 0.1kg/m^2^ on average, then a shift of just 5% in the allele frequency would increase average BMI in the population by 5kg/m^2^, i.e. from 25 to 30 (10).

Intergenerational GWAS studies have identified rapid gene pool changes in humans that have contributed to changes in phenotype (11). Global myopia rates have doubled over the past three decades (12). Twin studies find myopia highly heritable, at 50%-65% (13), GWAS studies have identified myopia-prone genes that explain between 35% and 40% of myopia heritability (13). A recent GWAS study (11) found in the UK that since 1940 a population-wide increase in multiple myopia alleles has occurred and explains a small, but significant, proportion of the UK myopia epidemic. Given it is plausible that a rapid dramatic environmental change can cause a rapid gene-pool change for traits with large standing genetic variation, such as obesity, next we explore if the change presented by modern obstetrics was dramatic enough to result in a rapid change to the gene pool.

### The Obstetrical Miracle Changed the Evolutionary Pressures on Obesity

The high risk of death during childbirth played a large role in shaping human evolution. However, with modern obstetrics, developed nations today have maternal death rates (MDR, share of women that will die from maternal causes in their lifetime) well below 1%, that no longer presents a strong evolutionary pressure in these nations, especially when combined with low total fertility rates being a result of very efficient birth control. Yet, as recently as 1990, several African nations had MDRs comparable to the pre-obstetric era, with multiple nations above 10%. In most developed nations, the introduction of novel obstetrical care between 1935 and 1945 led to a dramatic and rapid reduction in maternal mortality dropping over 20-fold in ten years (4) with MDRs no higher than a fraction of a percent. During the same period, perinatal mortality (neonate death in the first month of life) dropped precipitously from as high as 14% down to 0.4% (4).

Prior to modern obstetrics the predominant causes of maternal and perinatal mortality included: major hemorrhage, hypertensive disease, specifically pre-eclampsia and eclampsia, infection, asphyxia, and placental abruption (4). Obesity increases the risk of all the major historical causes of childbirth-related deaths dramatically (5). Even with modern obstetrics, obesity increases the relative risk of maternal and perinatal mortality by approximately 3-4 fold (5, 14, 15). Prior to modern obstetrics, obesity likely placed a similar if not greater relative risk on childbirth. Therefore in developed countries with an MDR of 0.01%, obese women would have an MDR of approximately 0.04%, this absolute difference in death rate would result in only a negligible evolutionary pressure. However, in the pre-obstetric period, obesity would present a high absolute risk, creating a strong negative selection pressure. Sierra Leone in 1990 had an MDR of nearly 15%; historically the MDR of all nations was likely as high if not higher. Consider a nation prior to modern obstetrics with an MDR of 15%. In such nations obese women would have an MDR of 60%, given the known relative risk obesity carries with childbirth is four times greater. In addition, prior to modern obstetrics perinatal mortality was approximately 14% of which obesity increases relative risk also by as much as 4 fold, suggesting prior to modern obstetrics an obese woman who survived childbirth, will have as many as 56% of their newborns die in the first month of life. Yet despite this large disproportionate loss of obesogenic genes due to childbirth deaths, the gene-pool remained stable, suggesting a stabilizing pressure existed.

The sudden removal of the negative selection pressure would have led to a rapid increase in obesity prone alleles. Multiple hypotheses have been proposed to identify the potential positive selection pressures on obesity. These have been discussed in detail elsewhere (16, 17). The microevolutionary hypothesis does not question their potential validity. In addition, a positive pressure for obesity must also have been a floor effect with too many alleles predisposing to low BMI being not sustainable with life, given a BMI of 11 is considered the lowest limit for survival (18). In addition, low BMI is a well recognized risk factor for infertility, with the risk increasing as BMI decreases (19). The floor effect alone represents a potential positive stabilizing selection pressure towards higher BMI, while childbirth represented a counter negative selection pressure.

Modern medicine has not removed the evolutionary pressure represented by the floor effect against low BMI. However, the sudden environmental change introduced by modern obstetrics since 1935-1945 did result in the exact conditions required for rapid gene-pool change for a trait like obesity with large standing genetic variation. Examining in detail the time course of the obesity epidemic can elucidate if this matches when the phenotype changes would be expected to be observed following the introduction of obstetrics.

### The Timing and Pattern of the Obesity Epidemic

The obesity epidemic is thought to have begun in the 1970s-1980s (20). However, when examining the timing of obesity by birth cohorts, Komlos and Brabec (21) found in the U.S. BMI began rising quickly in those born after 1945 directly after the introduction of modern obstetrics. This phenomenon has been referred to as an obesity cohort effect with generations born more recently having higher rates of obesity, when compared with older generations, and this has been observed in multiple developed nations (22–28). The population born after modern obstetrics first reached adulthood in the mid 1960s, but by 1970 they represented less than 10% of the US adult population (29) and while they did experience a higher rate of obesity than previous generations in the 1960s, the average adult obesity rate appeared to rise only slightly during this time given their relatively small percentage of the population. By 1980, this age segment grew to represent 50% of the population (29), becoming a significant percent of the adult population and with this the average adult obesity rate began to rise quickly, and the obesity epidemic began.

### Direct Evidence Supporting the Microevolutionary Hypothesis

The evidence presented so far, while not conclusive, does demonstrate that a rapid change in the obesogenic gene pool is plausible, and should be considered when interpreting obesity genetic studies. With this understanding, multiple studies examining obesity genetics reveal evidence supporting that a rapid change in the gene pool has occurred along with the timing expected by the microevolutionary hypothesis. Twin studies comparing obesity genetic variance by birth cohort consistently identify the additive genetic variance of BMI is increasing in those born after the introduction of modern obstetrics with limited changes in environmental variance (30–32). Without considering the microevolutionary hypothesis, these consistent findings appeared counter-intuitive given most proposed causes of the obesity epidemic have been believed to be environmental changes, well explained by Reddon *et al*. (32). *“One would presume the emergence of a society characterized by food abundance and physical inactivity may increase the impact of environment (and therefore decrease the impact of genes) in the determination of the obese phenotypes. Counter intuitively, the proportion of variability in BMI attributable to genetic variation is increased among people born after the establishment of a modern ‘obesogenic’ environment*.*”* This finding is well demonstrated by a Swedish twin study, in which they found additive genetic variance of BMI increased significantly between Swedish twin cohorts born in 1951 versus 1981 from 4.3 to 7.9 (31). This increasing genetic variance in those born in later birth cohorts was significantly correlated with the increasing obesity rate of the population, while the environmental variance of BMI showed no association with the population obesity rate. This pattern of increased genetic variance for BMI, with limited change in environmental variance is exactly what the microevolutionary hypothesis would predict, a population with wider variation of obesity promoting genes. However, these findings were interpreted presuming a change in the obesogenic gene pool is not possible, and suggested the only reasonable way to explain these results is a gene x environment interaction. While it is true these findings could be explained by a gene and environment interaction (and this also may be occurring), a more parsimonious explanation of these results is that a change in the gene pool has occurred. If environmental change increased BMI via a gene and environment interaction one would predict both genetic variance and environmental variance to be increased, not only heritability.

In addition, several longitudinal GWAS studies have identified direct evidence of increases in several of the obesogenic alleles in generations born after the introduction of modern obstetrics. Yet the hidden heritability of obesity and GWAS can only explain a limited percent of the known obesity heritability. Therefore, modern GWAS studies can not demonstrate if these changes are or are not producing a measurable effect on the obesity epidemic. However, the pattern of increasing obesogenic genes does support the microevolutionary hypothesis by itself. For example Rosenquist *et al*. (33) reported the gene pool frequency of the well known obesity promoting allele FTO pre and post 1942. Comparing the frequency of the obesity prone homozygote (AA) and heterozygote (AT), with the frequency of the obesity protective homozygote (TT) in the birth cohorts born before 1942 finds the pre 1942 frequency of the AA/AT was 63.1% compared with the post 1942 frequency of 66.7%. This increase in frequency of the obesity prone homozygote and heterozygote is statistically significant (Chi squared = 24.17 P<0.0001) (For calculation summary see supplementary data.) In addition Walter *et al*. (34) examined genetic risk score with BMI across different birth cohorts. They reported allele frequency across 4 birth cohorts in those born pre 1924 through those born as late as 1958. Of the 29 SNPs they examined, approximately 10% of the obesity prone alleles significantly increased in frequency in the later birth cohorts among population studied, with no obesogenic SNPs decreasing in frequency.

Budnik and Henneberg (35), found that greater national obesity rates are associated with reduced opportunity for natural selection over the past century. Given their findings they proposed that a portion of the obesity epidemic may have been caused by a reduction in evolutionary pressure leading to a change in the gene pool. However, they did not identify a mechanism for how reduced natural selection could increase obeseogenic alleles in the population. A reduction in maternal and perinatal mortality does offer a plausible mechanism explaining why reduced natural selection would be associated with increasing obesity rates. A recent GWAS study by Wu *et al*. (36) examined obesity genetic risk score (combines known obesogeneic alleles to offer a risk of obesity for an individual), across birth cohorts and infant mortality rate during birth year. The study found the number of obesogenic alleles increasing in cohorts born since the 1930s, and in addition, found this increase in obesogenic alleles correlated with infant mortality. The findings of Wu et al (36) offer evidence suggesting that natural selection has occurred in contemporary humans with childbirth related mortality acting as a selection pressure on BMI, exactly as would be predicted by the microevolutionary hypothesis.

A change in the obesogenic gene pool is not only possible but evidence suggests it may have occurred, the wide-spread belief that the obesity epidemic could not have been caused by a rapid change in the population gene-pool can no longer be presumed. Alternatively despite evidence supporting a rapid change in the gene pool, this genetic evidence alone only demonstrates plausibility, and is not sufficient to demonstrate, changes in the gene pool are the major driving force of the obesity epidemic. While the rise of BMI in any nation is certainly multi-factorial, with different obesity drivers having larger influences in various nations, examining how the microevolutionary hypothesis can offer an explanation for the global nature of the obesity epidemic offers a parsimonous mechanism tying together the global trends of increasing BMI across disparate nations.

### The Microevolutionary Hypothesis: A Parsimonious Explanation for the Obesity Epidemic

Many factors have been identified to influence obesity such as calorie intake and physical activity in addition to microevolutionary change. Importantly, the microevolutionary hypothesis does not challenge that these factors can also have an influence on obesity. However, these and other proposed factors have been insufficient to explain the global nature of the obesity epidemic. This is why there remains no scientific consensus on the cause of the obesity epidemic (37).

Between 1990 and 2016, obesity rates in every nation increased (ourworldofdata.com). During this same time period, approximately 25% of nations had decreased or stable total calorie intakes (Our world of data). During this time in these nations with stable or decreased calorie intakes, the median increase in obesity rate was approximately 10%. Conversely, examples exist of nations prior to the obesity epidemic in which calorie intake increased with no effect on obesity rate. For example, Switzerland increased calorie intake 17% from ∼ 3,000 kcal per day in 1947 to 3,500 kcal per day in 1961 with a negligible change in obesity rate. Switzerland’s calorie intake has oscillated up and down between 3,300 and 3,500 kcal per day between 1975 and 2016 while obesity rates consistently increased from 5% to 20% during this period of relatively unchanged calorie intake (Our world of data). This same pattern of increasing calorie intake prior to the obesity epidemic and stable calorie intake during the obesity epidemic has been observed in several other nations including Germany, Poland, France and Finland (our world of data). In addition, multiple nations have been identified to have increasing obesity rates despite increasing average levels of physical activity (38–40).

The microevolutionary hypothesis helps to explain these counter-intuitive discrepancies, as nations with high childbirth related deaths would have strong stabilizing evolutionary pressure keeping the obesity gene pool constant in these populations (ie, increasing calories would have little effect on obesity given the relatively low number of obesity prone alleles including alleles with gene and environment interactions). Conversely, after modern obstetrics is introduced nations would begin to experience an increase in obesogenic genes which are both environmentally dependent and independent. After the introduction of modern obstetrics the microevolutionary hypothesis would predict obesity rates will rise with or without increasing calorie intake or physical inactivity, but would likely rise faster with increased calorie intake or physical inactivity. However, prior to modern obstetrics calorie intake and physical inactivity rates would have little influence on obesity rates, given the relatively small percent of obesogenic alleles in the population, and of these alleles only a fraction would be obese due to gene and environment interactions.

### Hypothesis Predictions

Some of the predictions of the microevolutionary hypothesis are difficult to test directly because of the hidden heritability of obesity, and the limitation of GWAS studies. Yet the hypothesis does make several predictions that are directly testable using existing data. Specifically, a nation’s earlier maternal death rate (MDR) should be inversely associated with a nation’s obesity rate today. Earlier MDR should be able to offer additional explanations of obesity rate over confounding variables known to be associated with increasing obesity rates. The data on MDR for all nations globally were consistently collected since the year 1990 and varied from 0.01% to 14.97%, with a median of 0.48%. The hypothesis would predict that comparing only nations with MDR below the median in which all nations have MDRs that are less than a fraction of a percent, the MDR should have no association with obesity rates given the variation among MDRs in these nations would produce a negligible difference between these nations in terms of evolutionary pressure. Alternatively, nations with MDR above the median offer a wide range of MDR values (0.48%-14.97%) placing variable evolutionary pressures and in these nations MDR should be more predictive of obesity rates. Considering the example of Switzerland showing calorie intake changes had no association with obesity, prior to 1961, the hypothesis would predict in nations with MDR above the median that calorie intake and physical inactivity will have limited association with obesity rate given the relatively lower frequency of environmentally dependent obesogenic alleles in the population. However, calorie intake and physical inactivity are expected to be associated with obesity rate in the nations with MDR below the median given the frequency of environmentally dependent obesogenic alleles is a higher relative percent of these alleles in the population. A third prediction is that no nation with high maternal mortality rates will have a high obesity rate.

## Results

Figure 1 shows regressions of obesity prevalence rates in 1990 and in 2016 on MDR in 1990 . Regressions are of the same kind – exponential and of the same strength, about 50% of variance in obesity rate in each case is explained by MDR.

**Figure 1.**
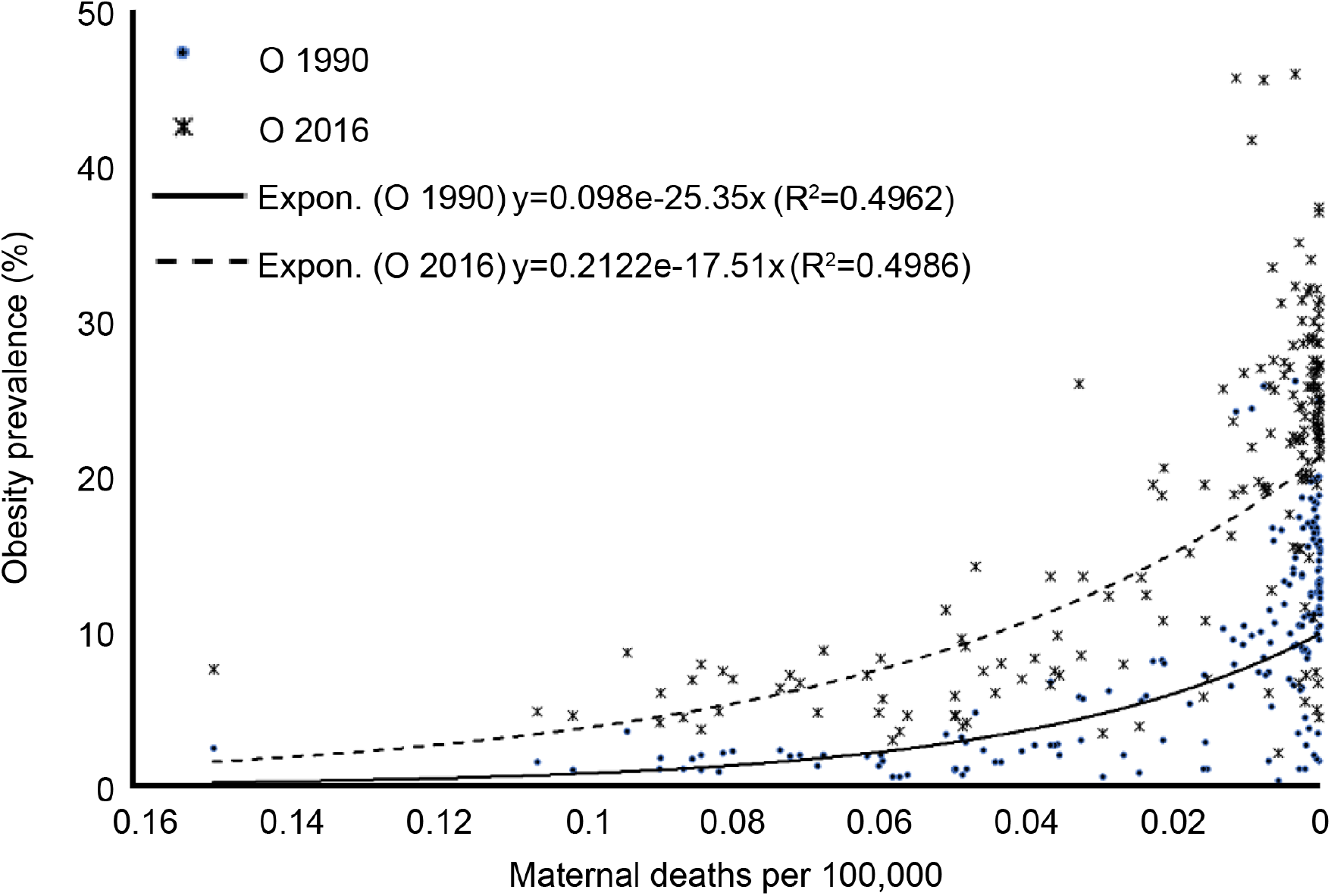
Regressions of obesity rates per country in 1990 and in 2016 on the rate of maternal deaths in the same year. Please note the strong negative relationship R=-0.70 -- the less maternal deaths, the greater the obesity prevalence. Strength of the relationship is the same for 1990 as for 2016.

Obesity rates among countries in 1990 and in 2016 were strongly correlated (Spearman’s “rho” = 0.955, Pearson’s moment product r=0.969, p<0.001, N=191), thus further analyses were conducted using only the 2016 obesity rates.

Maternal death rates of 191 countries correlate strongly, negatively and significantly with those countries’ obesity prevalence rates (Table 1). They also correlate significantly with GDP, physical inactivity, caloric consumption and antibiotic use. These variables, that also correlate with obesity rates, were considered confounders and thus kept statistically constant when exploring the correlation between MDR and obesity in partial correlation analysis finding a moment-product r of -0.336 (p=<0.001, N=133) (Table S1).

**Table 1.**
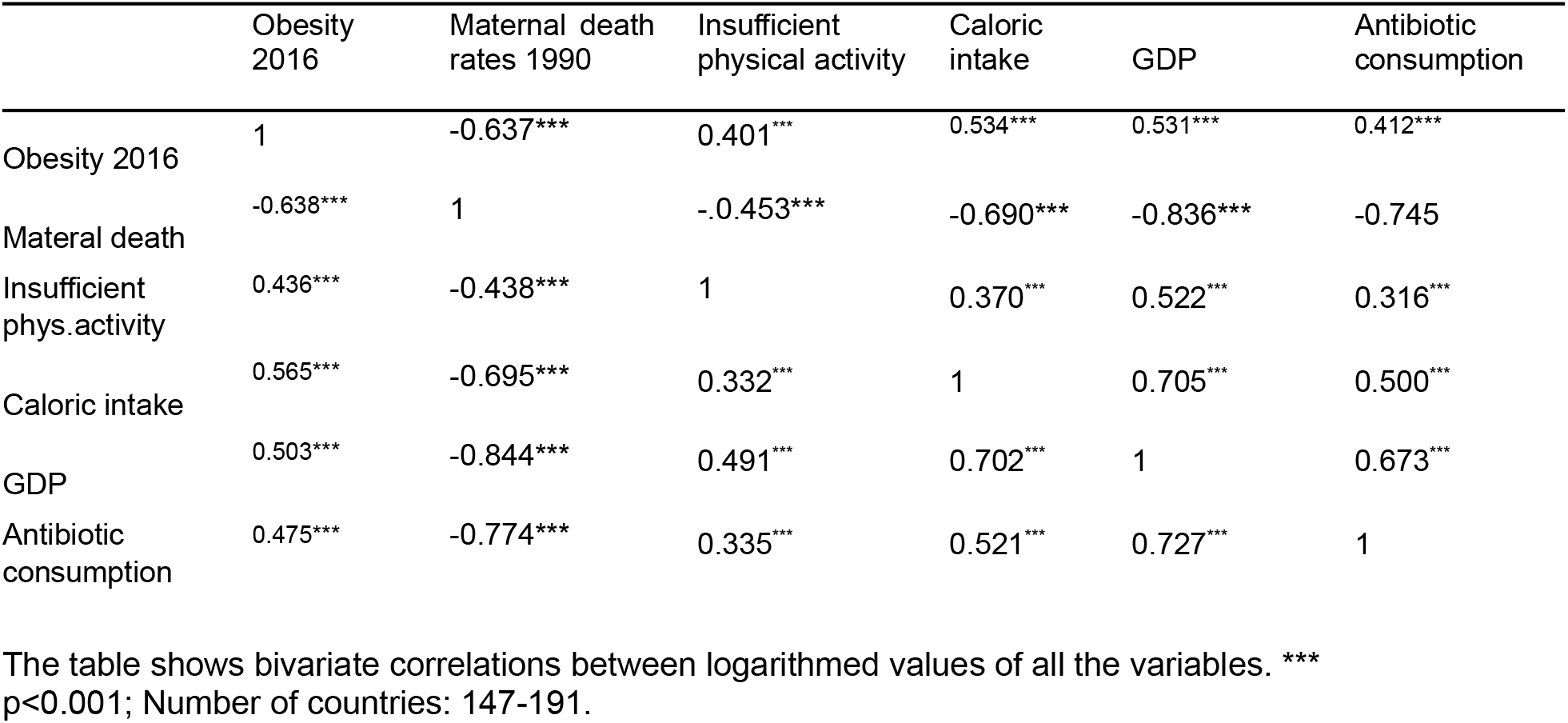
Pearson r (above the diagonal) and nonparametric Spearman’s rho (below the diagonal) for all variables studied.

When logarithmed obesity prevalence rates are treated as a dependent variable and independent variables are logarithmed MDR, GDP, calorie intake, insufficient physical activity and antibiotic consumption in the linear multivariate regression analysis (Table 2), MDR was the most significant of the studied variables. Insufficient physical activity and caloric intake were, as expected, significant, but had lower beta coefficients indicating that their effect size on obesity was less strong than that of MDR. In a stepwise multivariate regression analysis using probability of F to enter <=0.05 and probability of F to remove >= 0.10 only maternal death rates 1990 and caloric intake 2017 were entered, while the other three confounders were removed. With 2016 obesity rates held as the dependent variable MDR showed a β coefficient of -0.179 (SE=0.031, R^2^= 0.406 p<0.001) while caloric intake showed a β coefficient of 0.789, (SE=0.395, R^2^= 0.423, p<0.047 (Table S2).

**Table 2.**
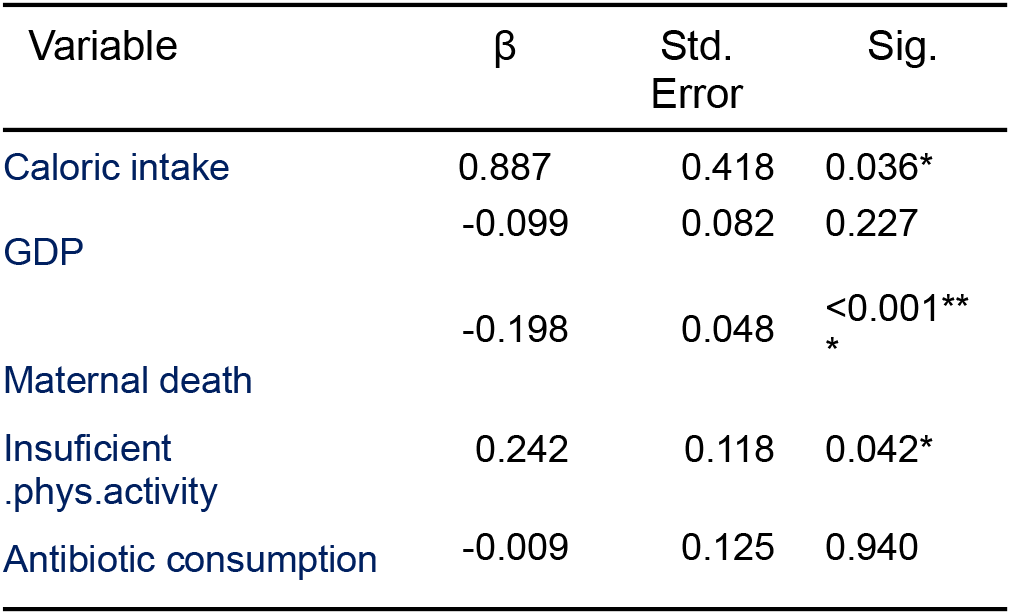
Independent predictors of obesity prevalence rate 2016 based on multiple linear regression modelling using logarithmed variable values. Df1=5, df2=133. Adjusted R^2^ = 0.422.

| Variable | $\beta$ | Std. Error | Sig. |
| --- | --- | --- | --- |
| Caloric intake | 0.887 | 0.418 | 0.036* |
| GDP | -0.099 | 0.082 | 0.227 |
| Maternal death | -0.198 | 0.048 | <0.001**<br>* |
| Insufficient<br>.phys.activity | 0.242 | 0.118 | 0.042* |
| Antibiotic consumption | -0.009 | 0.125 | 0.940 |

The median maternal death rate in 1990 was 0.48%. We explored relationships among MDR, obesity and the confounding variables separately in nations above and below this median. Nations with a MDR below the median showed a range of MDR of <0.01% to 0.45%. Nations with a MDR above the median showed a range of MDR of 0.48% to 14.93%. Results for countries above the median MDR, shown in Tables 3 and 4 indicate a stronger association compared to all nations relationship between MDR and obesity. In nations with MDR above the median all variables showed a significant association with obesity rates and MDR (Table 3) suggesting all other variables are potential confounders. When the four confounding variables are held constant to perform a partial correlation analysis between MDR and 2016 obesity rates in nations with above the median MDR a coefficient of r=-0.573 (p<0.00001) is obtained (Table S3). This association between MDR and obesity in nations with MDR above the median when confounders are held constant demonstrates a significantly stronger relationship than the one found across all countries (comparing r=-0.573 with r=-0.336 yields a significant difference p=0.03, when tested by (https://www.psychometrica.de/correlation.html). In multiple regression analysis for this group of countries, with MDR above the median the MDR turned out to be the only independent variable significantly influencing obesity rate,with GDP, calorie intake, physical inactivity and antibitoic use showing no independent association with obesity rate in these nations (Table 4, Figure 2 and Figure S1).

**Table 3.** Pearson r (above the diagonal) and nonparametric Spearman’s rho (below the diagonal) correlation between all variables studied in a sample of countries with maternal death rate above the median (>=0.45 per 100,000).

|  | Obesity<br>2016 | Maternal death<br>rates 1990 | Insufficient<br>physical activity | Caloric<br>intake | GDP | Antibiotic<br>consumption |
| --- | --- | --- | --- | --- | --- | --- |
| Obesity 2016 | 1 | -0.705*** | 0.285<sup>*</sup> | 0.356** | 0.428*** | 0.306** |
| Maternal death | -0.695*** | 1 | -.0.373** | -0.457*** | -0.620*** | -0.468*** |
| Insufficient<br>phys.activity | 0.334** | -0.385*** | 1 | 0.250<sup>*</sup> | 0.408*** | 0.165 |
| Caloric intake | 0.319** | -0.438*** | 0.248<sup>*</sup> | 1 | 0.450*** | 0.105 |
| GDP | 0.449** | -0.647*** | 0.405** | 0.418** | 1 | 0.438*** |
| Antibiotic<br>consumption | 0.330** | -0.567*** | 0.186 | 0.067 | 0.540*** | 1 |

The table shows bivariate correlations between logarithmed values of all the variables. \*\*\* p<0.001; Number of countries: 64-90.

**Table 4.**
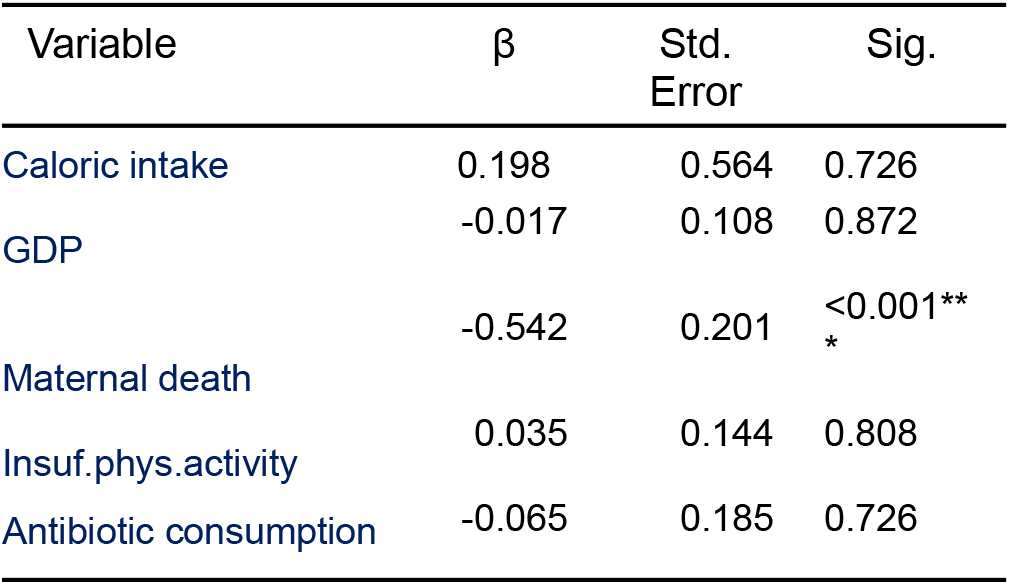
Independent predictors of obesity prevalence rate 2016 in countries with greater than median maternal death rate (>= 0.45 per 100,000) based on multiple linear regression modelling using logarithmed variable values. Df1=5, df2=58. Adjusted R^2^ = 0.457.

| Variable | $\beta$ | Std. Error | Sig. |
| --- | --- | --- | --- |
| Caloric intake | 0.198 | 0.564 | 0.726 |
| GDP | -0.017 | 0.108 | 0.872 |
| Maternal death | -0.542 | 0.201 | <0.001**<br>* |
| Insuf.phys.activity | 0.035 | 0.144 | 0.808 |
| Antibiotic consumption | -0.065 | 0.185 | 0.726 |

**Figure 2.**
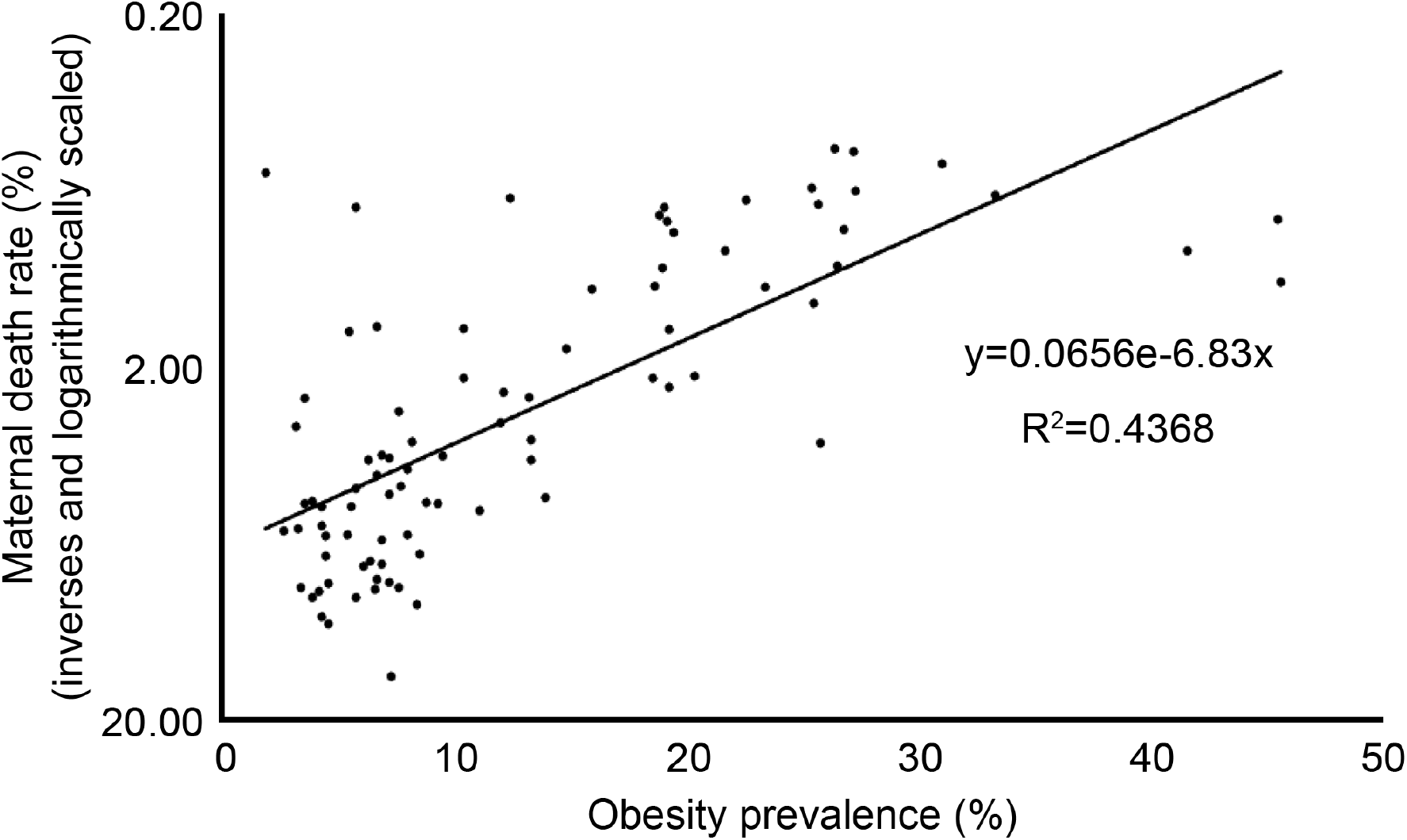
Relationship between the obesity prevalence rates and maternal death rates in countries with maternal death rates above the median value.

In the group of countries with lower than median MDR, obesity rate did not correlate significantly with MDR (r=-0.067, rho=0.107, N=90, p=0.863) and it did not show significant contribution to obesity variance in multiple regression analyses. In this group of nations, with MDR below the median, caloric intake and insufficient physical activity do show significant association with obesity rate (Table S4).

Additional analyses were performed only in developing nations to identify the frequency of nations with both high obesity rates and a high Maternal Mortality Rate (MMR). While no clear globally accepted definition of a high obesity nation exists, the definition was obtained using the 2017 obesity update of the Organization for Economic Cooperation and Development (OECD), which reported across the OECD nations 19.5% of the adult population was obese in 2015 (41). Given a national obesity rate of 19.5% is below the global median rate, it represents a conservative threshold to define a “high obesity nation”, including more than half the nations of the world.. Nations with obesity rates higher or lower than 19.5% were referred to as high or low obesity rate nations. High MMR was defined as above 500 deaths in 100,000 live births, given this would include nations with MMRs the World Health Organization classifies as very high or extremely high. Nations with MMRs above or below 500 deaths in 100,000 live births were referred to as high or low MMR nations respectively. Developing nations were defined using those listed as such by the United Nations World Economic Situation and Prospects (WESP).

Given these definitions 128 developing nations were identified with data on obesity rate and MMR. Of these nations 80 were found to have a low MMR and 58 (72.5 %) of these low MMR nations had high obesity rates. Among 48 developing nations with high MMR, only one nation had a high obesity rate (2%). The difference in high obesity rates between nations with low MMR (72..5%) and high MMR (2%) is large and significant (Chi Squared=42.592 P<0.00001). To ensure these results were not entirely driven by the more developed nations, the same analysis was performed in only the least developed nations, and only in low-income nations as defined by WESP. All analyses identified multiple nations with low MMR to have high obesity rates, and only one nation with high MMR had high obesity rates all showing significant differences, details of these additional analyses can be found in the Supplementary Information.

## Discussion

The results of our three analyses come together to offer support of the microevolutionary hypothesis. Our primary analysis identified 1990 MDR to be strongly associated with modern obesity rates. With confounding variables held constant, 1990 MDR was demonstrated to be independently associated with national obesity rates explaining 11% of the variance between nations, while the other variables examined: GDP, calorie intake, physical inactivity, and antibiotic use, were all correlated with obesity variance, MDR was the most strongly correlated, with the largest effect size.

Our second analysis further supported the microevolutionary hypothesis. In this analysis the nations were divided in half with the median MDRs (0.48%) serving as the dividing line, examining nations above and below the median separately. In nations with MDR below the median no independent association of MDR with obesity rate was found after controlling for the other variable. This finding was what the microevolutionary hypothesis would predict, given the median MDR was 0.48%, and therefore when comparing nations below the median, the differences in MDR are a fraction of a percent and therefore would produce negligible differences in negative selection pressure to be able to identify differences in obesity rates over such a short time scale. However when comparing nations with MDR above the median the difference ranged from 0.48% to 14.93%, which creates large variation in negative selection pressure. As predicted in nations with high MDR, when confounding variables were controlled for, maternal death rate explains 33% of the variance in obesity rates between nations.

In addition this analysis found calorie intake and physical inactivity to be correlated with obesity in nations with MDR below the median, as would be expected. However, in nations with above median MDR, calorie intake and physical inactivity showed no association with obesity rate. This surprising finding that calorie intake and physical inactivity have no association with obesity rates in half of nations with MDR above the median calls into question if these two factors can be major drivers of the global obesity epidemic. Simultaneously it is logical why many obesity researchers concluded that the obesity epidemic’s major drivers are increased calorie intake and physical inactivity, given most studies on obesity have been conducted only in developed nations with low MDR, and in these nations calorie intake and physical inactivity are associated with obesity (42, 43). Yet given all nations have experienced an increasing obesity rate since 1990, we suggest that it is simply not possible for calorie intake and physical inactivity to be major drivers of the obesity epidemic if they show no association with obesity rate in half of the nations of the world. Interestingly, calorie intake and physical inactivity both have a similar range of values in nations above and below median MDR, with average calorie intake ranging from approximately 2,000 to 3,500 in both. Physical inactivity also showed a similar range in nations with above and below median MDR ranging from approximately 10% to 55% in both. The major driver of the obesity epidemic must explain without exception the global nature of the phenomenon. Obstetrical outcomes have improved in every nation globally without exception(Ourworldofdata.com), the microevolutionary hypothesis offers a parsimonious explanation for a single cause that has led to increasing obesity rates in all nations.

The third analysis performed only in developing nations found that those with low 1990 MMR, 72.5% have high modern obesity rates, but only one nation (2%), Haiti possessed a high MMR and high obesity rate. The definitions used to identify nations with high obesity and high maternal mortality were based on the WHO definition (>500 in 100,000 live births) and average obesity rates of OECD nations (19.5%), and were relatively arbitrary thresholds in their relationship to evolutionary pressures. Haiti’s MMR and obesity rate were both near these arbitrarily thresholds with a MMR of 625 in 100,000 live births and obesity rate of 22.7%. Interestingly the only other high MMR nations with obesity rates within 5% of our chosen threshold all had MMR rates near the MMR threshold as well suggesting evolutionary pressure is reduced at a higher MMR rate than the threshold chosen for the analysis.

### Study Limitations

The limitations of this study’s findings are that all analyses demonstrated statistical associations, of which causation can not be assumed given potential unforeseen confounders. It is possible some other medical interventions are the cause of the obesity epidemic and they may increase in use within a population at a similar temporal pattern as obstetrics lowered MDR.

Further limitations include the limited data for national MDR prior to 1990, and obesity rates prior to 1975 preventing closer examination of how earlier differences in MDR may have influenced obesity rates in different nations at earlier time points.

In addition, alternative hypotheses to a rapid gene pool change suggested by the microevolutionary hypothesis may explain the strong associations between MDR and obesity identified in this study. One such potential alternative hypothesis could be that heritability of obesity is explained by telomere length heritability rather than gene heritability. A second alternative hypothesis that could explain the MDR and obesity association is that maternal obesity influences long-term offspring obesity risk via in-utero maternal effects likely via altered hormone signals between obese mother and fetus.

### Social Implications

The hypotheses suggesting environmental changes in diet and physical activity as the major causes of the obesity epidemic, imply those with obesity must lack “willpower” to eat healthy and exercise. This wide-held belief has resulted in great social stigma for those suffering from obesity, indirectly blaming obese individuals for the cause of their state. While dieting and exercise can occasionally reduce body weight and thus BMI, nearly all long-term studies find dieting and exercise do not lead to sustained weight loss in the large majority of obese individuals. The microevolutionary hypothesis may alleviate some of the social stigma of obesity, by clarifying that the cause of most individuals’ obesity is the result of a collection of genes they were born with. Blame is not typically placed on those with genetic conditions that cause a pathological state. The microevolutionary hypothesis would hopefully help the general and medical communities understand that the cause of most individuals’ obesity is second to a genetic cause, and to stop stigmatizing obese individuals for the false belief that their lack of willpower is the cause of their obesity. With decreased stigma of obesity it is plausible both physicians and patients would be more open to treating obesity with effective medical interventions such as bariatric surgery and/or medications.

## Materials and Methods

### Data Extraction Methods

Using ourworldofdata.org, data were extracted for each nation for maternal mortality rate (MMR), and maternal death rate (MDR) (share of women that will die from maternal causes in their lifetime) at the earliest year available for which these data were available. For nearly all nations this year was 1990. For Obesity Rates data were obtained from 1990 and 2016, GDP per capita in 2019, calorie intake 2017. The national rate of insufficient physical activity was obtained from Guthold *et al*. (44), and antibiotic consumption in 2000 from Girum and Wasie (45).

### Data Analysis

First, scattergrams of relationships with curvilinear regressions fitted and non-parametric correlations (Spearman’s rho) of the variables studied were explored. Then natural logarithms of all data were calculated to bring their distributions closer to the homoscedasticity required for parametric correlations and regression analyses. Parametric correlation coefficients of log transformed variables were compared with non-parametric ones to ensure that parametric relationships were close to actual ones. Next, partial correlation analyses of Pearson moment-product correlations were carried out to stabilize the influence of confounding variables such as GDP, calorie intake and physical inactivity on the relationship between obesity and maternal death. Multivariate regression analyses, using obesity as the dependent variable were conducted to establish the strength and significance of the contribution of different variables to the variation in obesity. SPSS v 27 statistical package was used for these analyses.

The World Health Organization defines high Maternal Mortality Rate (MMR) nations when MMR is above 500 deaths in 100,000 live births, this definition was used for this analysis. MMR rates from 1990 were used, being the earliest year in which the MMR of nearly all nations was recorded. The definition of high obesity rate was obtained from the 2017 Obesity update of the Organization for Economic Cooperation and Development (OECD), which reported across the OECD nations 19.5% of the adult population was obese in 2015 (41). The obesity rate offered by the OECD nations was used as a conservative threshold to define nations with a high obesity rate as 19.5%. Obesity rates from 2016 were used as this was the most recent information available which included all nations globally.

The frequency of high obesity nations was compared between low MMR nations and high MMR nations. In an attempt to exclude the known association between developed nations and obesity, the analysis was performed in nations defined as developing nations by the United Nations World Economic Situation and Prospects report released in 2017 describing the nations the year prior in 2016 (46). The same analysis was performed in only low-income nations to help reduce the confounding effect of GDP on obesity. Low income nations included nations defined by the United Nations World Economic Situation and Prospects in 2017 as either low income or low middle income nations.

## Author Contributions

**Author 1:** Joseph Fraiman

Conceived and designed the analysis

Collected the data

Wrote the paper

-Wrote first draft and edited subsequent drafts and final version

Searched relevant literature

**Author 2:** Scott Baver

Wrote the paper

Edited subsequent draft and final version

Provided broad historical background

**Author 3:** Maciej Henneberg

Performed the analysis

Statistical analyses of all data, produced graphs and tables

Wrote the paper

Edited subsequent drafts and the final draft

Had multiple discussions with other authors

## Supporting Information

**Figure S1.**
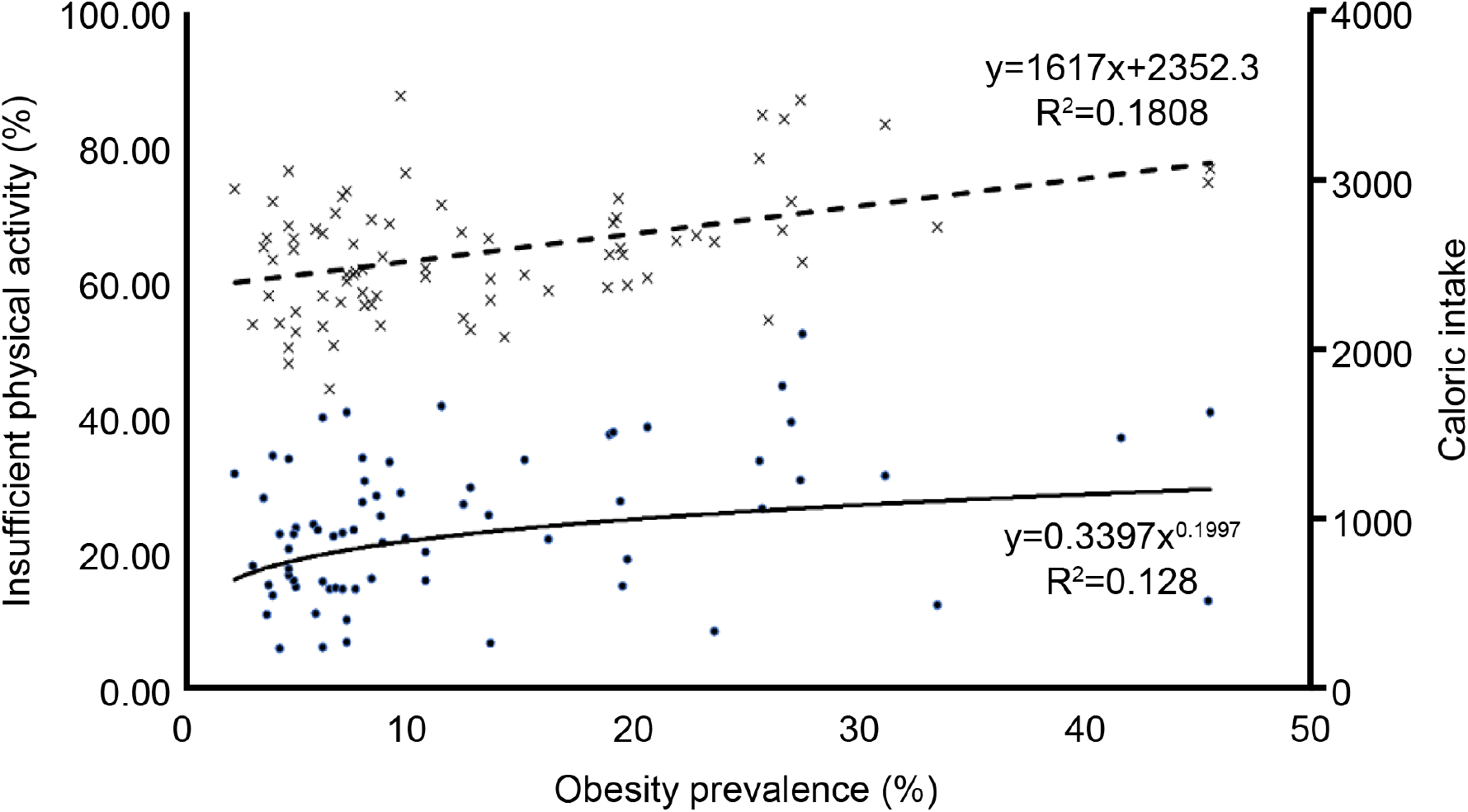
Relationship between the obesity prevalence rates and insufficient physical activity and caloric intake in countries with maternal death rates above the median value.

**Table S1.**
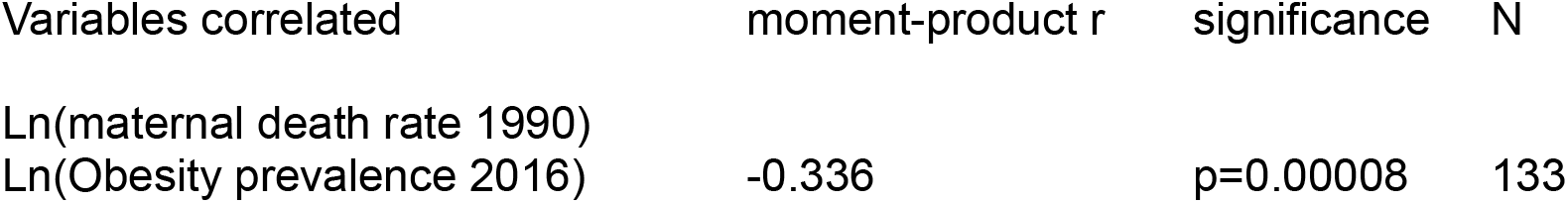
Result of the partial correlation analysis between maternal death rates 1990 and obesity rates 2016 when insufficient physical activity, caloric intake, GDP and antibiotic consumption are kept statistically constant.

**Table S2.**
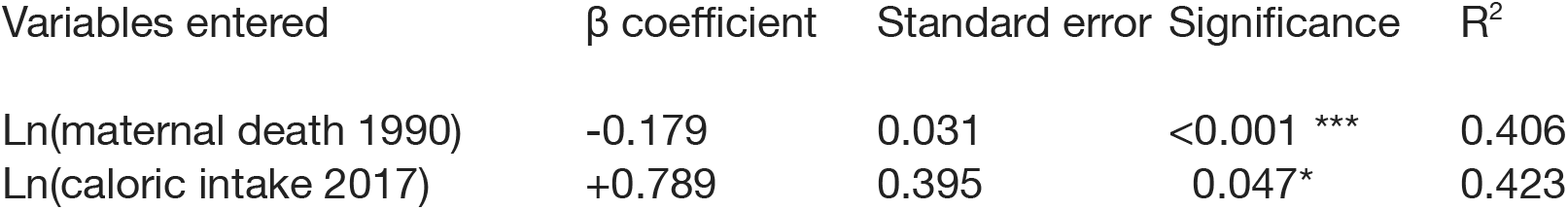
Results of the stepwise multivariate regression analysis using the obesity rates 2016 as the dependent variable.

**Table S3.**
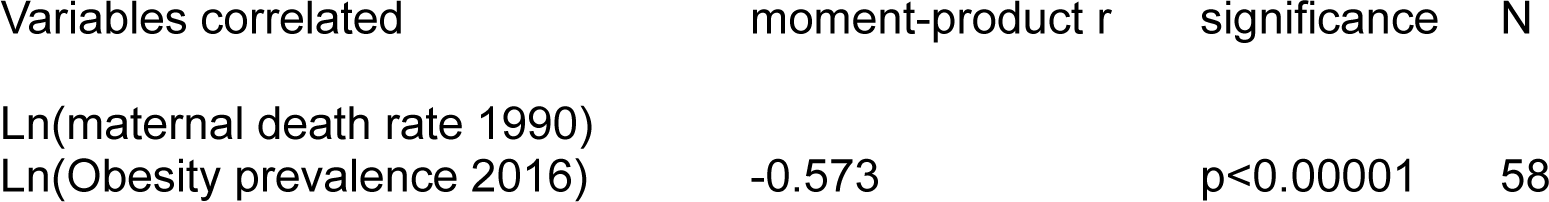
Result of the partial correlation analysis between maternal death rates 1990 and obesity rates 2016 in countries with greater than median MDR(>= 0.45 per 100,000) when insufficient physical activity, caloric intake, GDP and antibiotic consumption are kept statistically constant.

**Table S4.**
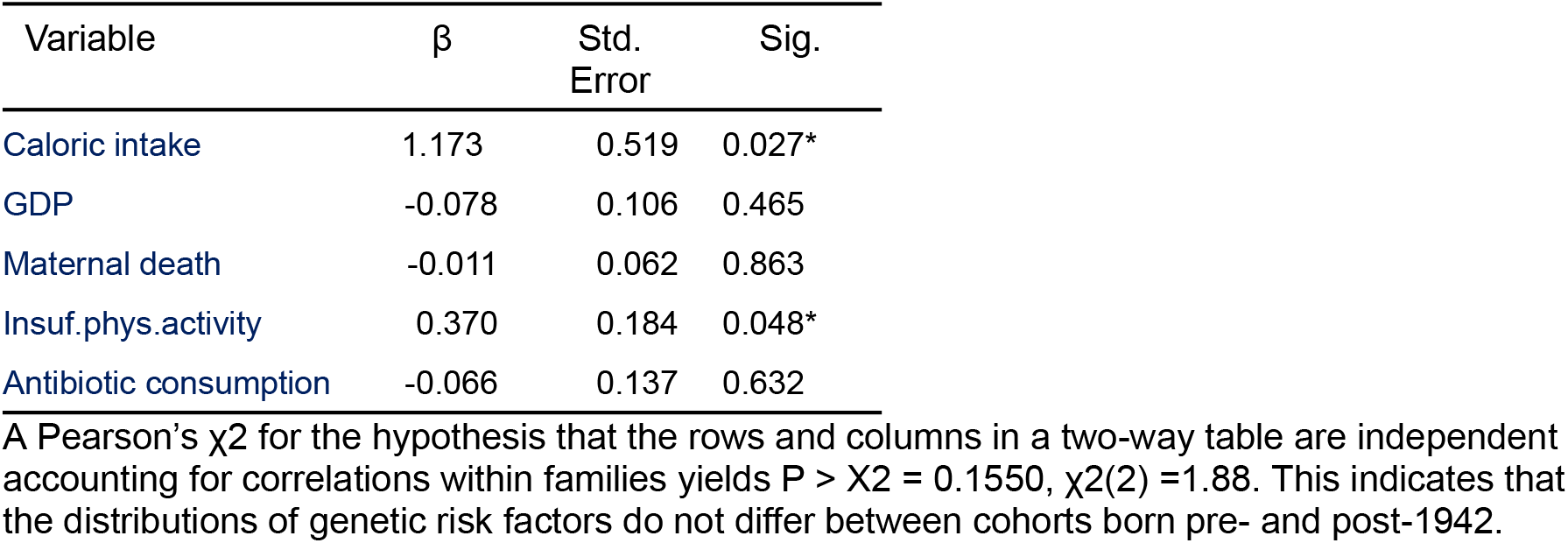
Independent predictors of obesity prevalence rate 2016 in countries with lower than median maternal death rate (>0.45 per 100,000) based on multiple linear regression modelling using logarithmed variable values. Df1=5, df2=65. Adjusted R^2^ = 0.120.

**Supplemental Analysis 1: FTO Allele Obesogenic Genotype Frequencies Pre and Post 1942**

FTO allele has been identified by GWAS studies to be associated with an increased risk of obesity, for both heterozygote (AT) and homozygote (AA), compared with homozygote (TT). The risk of obesity for AT was found to be an OR of 1.31 95% CI = 1.23 to 1.39), while for AA the OR was OR = 1.67; (95% CI = 1.47 to 1.89). With each additional allele adding 0.36 kg/m2, with homozygotes (AA) having higher BMI than heterozygotes (AT), and both having higher BMI than homozygote (TT) (1). The frequency of the obesity promoting FTO genotypes pre and post obstetrics were reported in Rosenquist *et al*. (2) using pre 1942 data and post 1942 data. The frequency of obesity promoting genotypes (AA/AT) were compared with obesity protective genotype (TT), prior to and after 1942. Rosenquist *et al*. (2) reported reported pre-1942 population frequency of obesity promoting genotypes (AA/AT) of 63.5% and obesity protective genotype (TT) of 36.5%. Post 1942 obesity promoting genotype AA/AT representing 66.9% of the population, while the obesity protecting genotype of TT represented 33.1%. This the change in frequency of the obesity promoting and protective genotypes pre 1942 and post 1942 is statistically significant (chi square X2=19.6232, P<0.00001).

**Supplemental Analysis 2: Additional Maternal Mortality Analyses In Least Developed and Lower Income Nations**

Examining only the least developed nations included a total of 47 nations, of which only 6 had low MMR, and 41 had high MMR. Of the least developed nations with low MMR three (50%) had high obesity, and of those with high MMR one (2.4%) had high obesity (Chi Squared=15.206 P<0.0001).

To ensure the results of these analysis were not mainly driven by differences in GDP between the developing nations a second exploratory analysis was performed only in nations of low and lower middle income. This analysis included 77 nations of which 36 were low income and 41 were lower middle income. Of the low and lower middle income nations 31 had low MMR and 46 had high MMR. High obesity was found in 17 (54.8%) nations with low MMR versus one (2.2%) nation with high MMR (Chi Squared=28.677 P<0.00001).

## References

1. C. E. Elks et al., Variability in the heritability of body mass index: A systematic review and meta-regression. Front. Endocrinol. 3, 29 (2012).

2. J. Belluz, Scientists Don’t Agree on What Causes Obesity, but They Know What Doesn’t (New York Times, New York, 2022).

3. R. L. Goldenberg, E. M. McClure, Maternal mortality. Am. J. Obstet. Gynecol. 205, 293–295 (2011).

4. R. L. Goldenberg, E. M. McClure, Maternal, fetal and neonatal mortality: Lessons learned from historical changes in high income countries and their potential application to low-income countries. Matern. Health Neonatol. Perinatol. 1, 3 (2015).

5. K. R. Andreasen, M. L. Andersen, A. L. Schantz, Obesity and pregnancy. Acta Obstet. Gynecol. Scand. 83, 1022–1029 (2004).

6. R. Barrett, D. Schluter, Adaptation from standing genetic variation. Trends Ecol. Evol. 23, 38–44 (2008).

7. T. Kunej et al., Obesity gene atlas in mammals. J. Genom. 1, 45–55 (2013).

8. G. Guo, H. Liu, L. Wang, H. Shen, W. Hu, The genome-wide influence on human BMI depends on physical activity, life course, and historical period. Demography 52, 1651–1670 (2015).

9. G. Gibson, Rare and common variants: Twenty arguments. Nat. Rev. Genet. 13, 135–145 (2012).

10. J. K. Pritchard, A. Di Rienzo, Adaptation – not by sweeps alone. Nat. Rev. Genet. 11, 665–667 (2010).

11. E. Long, J. Zhang, Natural selection contributes to the myopia epidemic. Natl. Sci. Rev. 8, nwaa175 (2021).

12. B.A Holden et al., Global prevalence of myopia and high myopia and temporal trends from 2000 through 2050. Ophthalmology 123: 1036–42. (2016)

13. R. L. Shah, J. A. Guggenheim, UK Biobank Eye and Vision Consortium, Genome-wide association studies for corneal and refractive astigmatism in UK Biobank demonstrate a shared role for myopia susceptibility loci. Hum. Genet. 137, 881–896 (2018).

14. M. Saucedo et al., Understanding maternal mortality in women with obesity and the role of care they receive: A national case-control study. Int. J. Obes. 45, 258–265 (2021).

15. Sebire et al., Maternal obesity and pregnancy outcome: a study of 287 213 pregnancies in London. International journal of obesity, 25, 1175–1182. (2001).

16. J. R. Speakman, Thrifty genes for obesity, an attractive but flawed idea, and an alternative perspective: The ‘drifty gene’ hypothesis. Int. J. Obes. 32, 1611–1617 (2008).

17. J. V. Neel, A. B. Weder, S. Julius, Type II diabetes, essential hypertension, and obesity as “syndromes of impaired genetic homeostasis” : The “thrifty genotype” hypothesis enters the 21st century. Perspect. Biol. Med. 42, 44–74 (1998).

18. James, W.P.T., Ferro-Luzzi, A. & Waterlow, j .C. Definition of chronic energy malnutrition in adults. Eur. f. Clin. Nutr. 42, 969-981 (1988).

19. M. Imterat, A. Agarwal, S. C. Esteves, J. Meyer, A. Harlev, Impact of body mass index on female fertility and ART outcomes. Panminerva Med. 61, 58–67 (2019).

20. Q. Zhang, Y. Wang, Trends in the association between obesity and socioeconomic status in U.S. adults: 1971 to 2000. Obes. Res. 12, 1622–1632 (2004).

21. J. Komlos, M. Brabec, The trend of BMI values of US adults by deciles, birth cohorts 1882–1986 stratified by gender and ethnicity. Econ. Hum. Biol. 9, 234–250 (2011).

22. M. A. Allman-Farinelli, T. Chey, A. E. Bauman, T. Gill, W. P. T. James, Age, period and birth cohort effects on prevalence of overweight and obesity in Australian adults from 1990 to 2000. Eur. J. Clin. Nutr. 62, 898–907 (2008).

23. O. K. Caman et al., Longitudinal age-and cohort trends in body mass index in Sweden – a 24-year follow-up study. BMC Public Health 13, 1–10 (2013).

24. I. Diouf et al., Evolution of obesity prevalence in France: An age-period-cohort analysis. Epidemiology 21, 360–365 (2010).

25. B. K. Jacobsen et al., Increase in weight in all birth cohorts in a general population: The Tromsø study. Arch. Intern. Med. 161, 466–472 (2001).

26. M. Lahti-Koski, P. Jousilahti, P. Pietinen, Secular trends in body mass index by birth cohort in eastern Finland from 1972 to 1997. Int. J. Obes. 25, 727–734 (2001).

27. L. W. Olsen, J. L. Baker, C. Holst, T. I. A. Sørensen, Birth cohort effect on the obesity epidemic in Denmark. Epidemiology 17, 292–295 (2006).

28. B. L. Thomsen, C. T. Ekstrøm, T. I. A. Sørensen, Development of the obesity epidemic in Denmark: Cohort, time and age effects among boys born 1930–1975. Int. J. Obes. 23, 693–701 (1999).

29. United Nations, United Nations World Economic Situation Prospects 2020 (United Nations, New York, 2020).

30. B. Rokholm et al., Increased genetic variance of BMI with a higher prevalence of obesity. PLoS One 6, e20816 (2011).

31. B. Rokholm et al., Increasing genetic variance of body mass index during the Swedish obesity epidemic. PLoS One 6, e27135 (2011).

32. H. Reddon, J.-L. Guéant, D. Meyre, The importance of gene–environment interactions in human obesity. Clin. Sci. 130, 1571–1597 (2016).

33. J. N. Rosenquist et al., Cohort of birth modifies the association between FTO genotype and BMI. Proc. Natl. Acad. Sci. U. S. A. 112, 354–359 (2015).

34. S. Walter, I. Mejía-Guevara, K. Estrada, S. Y. Liu, M. M. Glymour, Association of a genetic risk score with body mass index across different birth cohorts. JAMA 316, 63–69 (2016).

35. A. Budnik, M. Henneberg, Worldwide increase of obesity is related to the reduced opportunity for natural selection. PLoS One 12, e0170098 (2017).

36. Y. Wu et al., GWAS on birth year infant mortality rates provides new evidence of recent natural selection. Proc. Natl. Acad. Sci. U. S. A. 119, e2117312119 (2022).

37. S. E. Ross, J. I. Flynn, R. R. Pate, What is really causing the obesity epidemic? A review of reviews in children and adults. J. Sports Sci. 34, 1148–1153 (2016).

38. J. S. Schiller, T. C. Clarke, T. Norris, Early Release of Selected Estimates Based on Data from the January-September 2017 National Health Interview Survey (Center for Disease Control and Prevention, Atlanta, GA, 2017).

39. S. Graf, M. Cecchini, Current and Past Trends in Physical Activity in Four OECD Countries: Empirical Results from Time Use Surveys in Canada, France, Germany and the United States, OECD Health Working Papers, No. 112 (OECD Publishing, Paris, 2019).

40. A. O. Werneck et al., Time trends and inequalities of physical activity domains and sitting time in South America. J. Glob. Health 12, 04027 (2022).

41. M. G. S. Devaux, Y. Goryakin, M. Cecchini, H. Huber, F. Colombo, OECD Obesity Update 2017 (2017).

42. C. O. Stubbs, A. J. Lee, The obesity epidemic: Both energy intake and physical activity contribute. Med. J. Aust. 181, 489–491 (2004).

43. J. O. Hill, E. L. Melanson, Overview of the determinants of overweight and obesity: Current evidence and research issues. Med. Sci. Sports Exerc. 31, S515–SS521 (1999).

44. R. Guthold, G. A. Stevens, L. M. Riley, F. C. Bull, Worldwide trends in insufficient physical activity from 2001 to 2016: A pooled analysis of 358 population-based surveys with 1·9 million participants. Lancet Glob. Health 6, e1077–e1086 (2018).

45. T. Girum, A. Wasie, Correlates of maternal mortality in developing countries: An ecological study in 82 countries. Matern. Health Neonatol. Perinatol. 3, 1–6 (2017).

46. UN.ESCAP, UN.ECA, UN.ECE, UN.ESCWA, UN.ECLAC, World Economic Situation and Prospects 2017 (United Nations, New York, 2017).

## SI References

1. T. M. Frayling et al., A common variant in the FTO gene is associated with body mass index and predisposes to childhood and adult obesity. Science 316, 889–894 (2007).

2. J. N. Rosenquist et al., Cohort of birth modifies the association between FTO genotype and BMI. Proc. Natl. Acad. Sci. U. S. A. 112, 354–359 (2015).

