## Supplemental Material for "Microevolutionary Hypothesis of the Obesity Epidemic"

### Supplementary Material
### Microevolutionary hypothesis of the obesity epidemic

Joseph Fraiman, MD[1,*], Scott Baver[2], PhD, Maciej Henneberg PhD, DSc, Dr h.c.[3]

***Corresponding Author:** Joseph Fraiman


**This PDF file includes:**

Supporting text
Figure S1
Tables S1 to S4
Legends for Movies S1 to Sx
Legends for Datasets S1 to Sx
SI References

**Supporting Information Text**

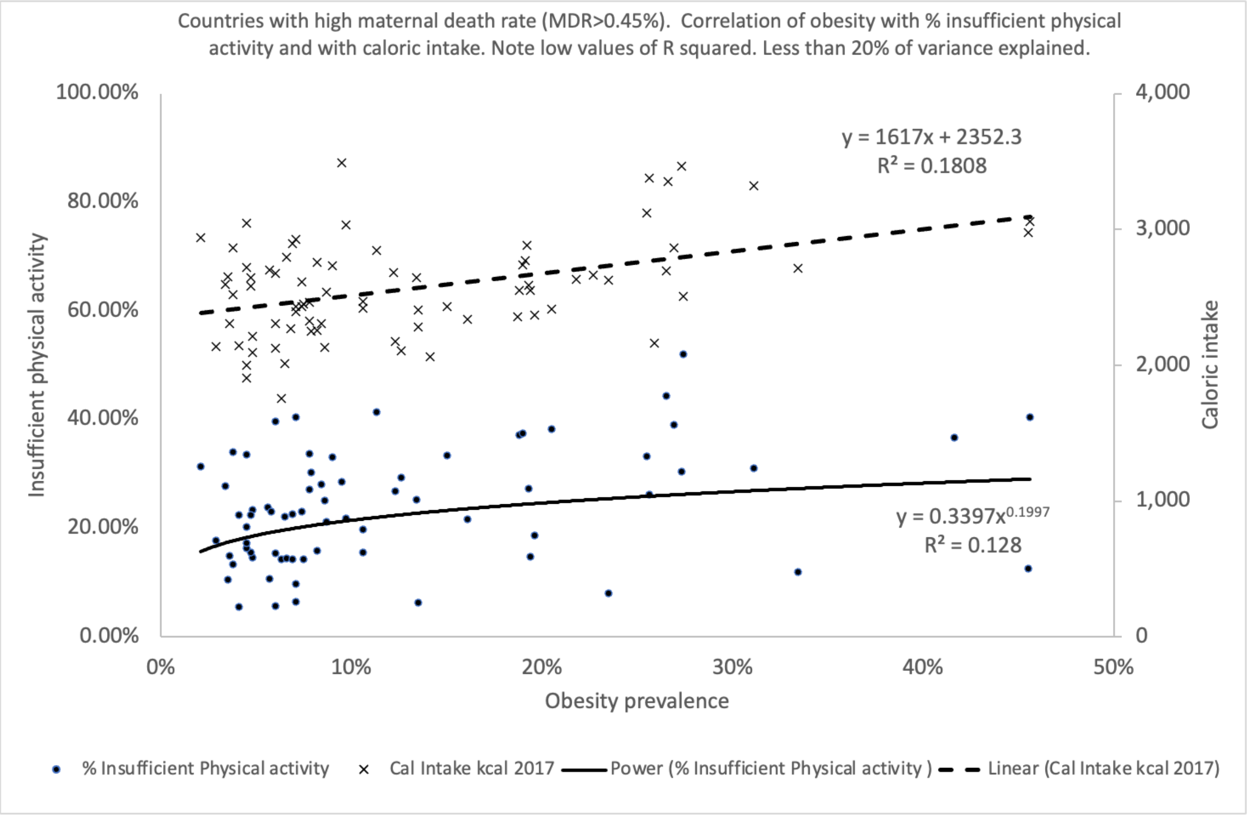


**Figure S1.** Relationship between the obesity prevalence rates and insufficient physical activity and caloric intake in countries with maternal death rates above the median value.

**Table S1.** Result of the partial correlation analysis between maternal death rates 1990 and obesity rates 2016 when insufficient physical activity, caloric intake, GDP and antibiotic consumption are kept statistically constant.

| Variables correlated | moment-product r | significance | N |
| --- | --- | --- | --- |
| Ln(maternal death rate 1990)<br>Ln(Obesity prevalence 2016) | -0.336 | p=0.00008 | 133 |

**Table S2.** Results of the stepwise multivariate regression analysis using the obesity rates 2016 as the dependent variable.

| Variables entered | β coefficient | Standard error | Significance | $R^2$ |
|---|---|---|---|---|
| Ln(maternal death 1990) | -0.179 | 0.031 | <0.001 *** | 0.406 |
| Ln(caloric intake 2017) | +0.789 | 0.395 | 0.047* | 0.423 |

**Table S3.** Result of the partial correlation analysis between maternal death rates 1990 and obesity rates 2016 in countries with greater than median MDR(>= 0.45 per 100,000) when insufficient physical activity, caloric intake, GDP and antibiotic consumption are kept statistically constant.

| Variables correlated | moment-product r | significance | N |
|---|---|---|---|
| Ln(maternal death rate 1990)<br>Ln(Obesity prevalence 2016) | -0.573 | $p<0.00001$ | 58 |

**Table S4.** Independent predictors of obesity prevalence rate 2016 in countries with lower than median maternal death rate (>0.45 per 100,000) based on multiple linear regression modelling using logarithmed variable values. Df1=5, df2=65. Adjusted $R^2$ = 0.120.

| Variable | β | Std. Error | Sig. |
|---|---|---|---|
| Caloric intake | 1.173 | 0.519 | 0.027* |
| GDP | -0.078 | 0.106 | 0.465 |
| Maternal death | -0.011 | 0.062 | 0.863 |
| Insuf.phys.activity | 0.370 | 0.184 | 0.048* |
| Antibiotic consumption | -0.066 | 0.137 | 0.632 |

A Pearson's χ2 for the hypothesis that the rows and columns in a two-way table are independent accounting for correlations within families yields P > X2 = 0.1550, χ2(2) =1.88. This indicates that the distributions of genetic risk factors do not differ between cohorts born pre- and post-1942.
